# Dissecting the Roles of the Tuberin Protein in the Subcellular Localization of the G2/M Cyclin, Cyclin B1

**DOI:** 10.1101/2021.10.25.465800

**Authors:** Jessica Dare-Shih, Adam Pillon, Jackie Fong, Elizabeth Fidalgo da Silva, Lisa A. Porter

## Abstract

Tuberin is a major component of the protein regulatory complex known as the Tuberous Sclerosis Complex and plays a crucial role in cell cycle progression and protein synthesis. Mutations in the Tuberin gene, *TSC2*, lead to the formation of benign tumors in many organ systems and causes the Tuberous Sclerosis Complex disorder. Genotypes ranging from point mutations to large deletions in the *TSC2* gene have been clinically characterized with a wide range of phenotypes from skin tumors to large brain tumors. Our current work investigates the molecular mechanisms behind Tuberin and its ability to regulate the cell cycle through its binding to the G2/M cyclin, Cyclin B1. After creating an early stop codon in a critical region of the Tuberin, our results show the *in vitro* phenotype that occurs from a truncated Tuberin protein. Herein we demonstrate that this clinically relevant truncated form of Tuberin promotes an increased nuclear accumulation of Cyclin B1 and a subsequent increase in cell proliferation supporting the phenotypic data seen in the clinic with Tuberous Sclerosis Complex patients showing deletions within the *TSC2* gene. This data provides an insight into some of the functional and molecular consequences of truncated proteins that are seen in clinical patients.

## Introduction

The Tuberin protein is encoded by the *TSC2* gene located on chromosome 16p13.3 consisting of 41 exons. It is comprised of 1804 amino acids and results in a 200 kDa protein [1]. This large protein contains a variety of conserved domains responsible for its function: two coiled-coil domains (aa346-371), a leucine zipper (aa75-107), a GTPase activating protein (GAP) homology domain (aa1517-1674) and a calmodulin domain (aa1740-1758) [2] (Fig. 1A). It is well-established that signals from the environment, including nutrient status, can alter the phosphorylation, protein-protein interactions and subcellular localization of Tuberin which are integrated to impact aspects of cell physiology [3–5]. The most well-known effector of Tuberin being the regulation of cell growth and protein synthesis through the downstream inhibition of the Mammalian Target of Rapamycin (mTOR) [5,6]. The Tuberin GAP domain inactivates mTOR signalling by stimulating auto-hydrolysis of Rheb-GTP, converting it to Rheb-GDP [4].

**Fig 1.**
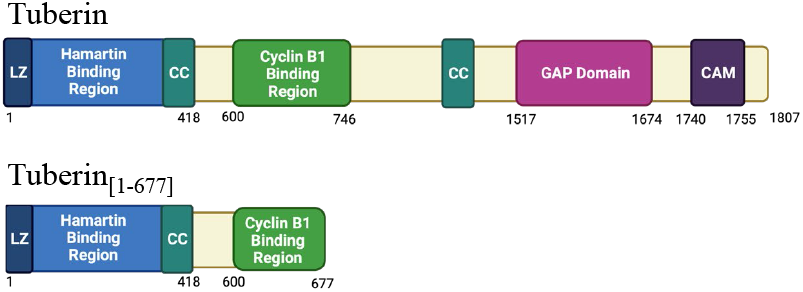
Simplified protein structure diagram of the Tuberin protein and Tuberin_[1-677]._ The primary structure of Tuberin (Top) and Tuberin_[1-677]_ (bottom) showing domains and binding patterns. The numbers below the diagram are the amino acids where each domain region starts and ends. LZ (Leucine zipper), CC (Coiled-coiled domain) and CAM (Calmodulin binding domain) are depicted. Images created with BioRender.com.

In addition to these roles, Tuberin also directly regulates the cell cycle, inhibiting progression through G1/S and G2/M under select environmental conditions [7–9]. During G1 phase of the cell cycle Tuberin, independent of Hamartin, interacts with the Cyclin-Dependent Kinase (CDK) inhibitor, p27, protecting p27 from ubiquitin-mediated degradation and permitting the nuclear accumulation of p27 by interfering with the 14-3-3b dependent cytoplasmic retention of p27 [7]. Through this mechanism, Tuberin markedly inhibits the G1/S transition [10]. In addition to inhibitory roles at the G1/S transition, Tuberin forms a transient complex with Cyclin B1 (CycB1) at the G2/M transition, providing a novel mechanism which CycB1 localization and mitotic onset is regulated [8]. Tuberin-CycB1 complex formation is in part regulated by the phosphorylation on 5 serine sites within the Cytoplasm Retention Signal (CRS) region of CycB1[8]. Tuberin has lower affinity to CycB1 when these serine sites are mutated to a glutamic acid (E) to mimic phosphorylation (CycB1-5E), and a higher affinity for binding CycB1 where these sites are altered to non-phosphorylatable alanines (A) (CycB1-5A) [8]. The formation of the Tuberin-CycB1 complex is dependent on serum levels, whereby ample nutrients permits complex formation that subsequently supports a delay in mitotic progression and an increase in cell size [9]. This work also shows that the Tuberin-CycB1 complex associates with both Hamartin and the G2 CDK, CDK1. Interestingly, CDK1 has been shown to phosphorylate Hamartin in the Tuberin binding domain to aid in the inhibition of pS6Kinase during the cell cycle [11]. Collectively, these data present the possibility that feedback from the Tuberin-CycB1 complex can be provided to regulate G1/S. Hence, Tuberin is a key integrator, sensing levels of nutrients, growth factors, and energy stores to regulate cell growth and cell proliferation [12,13].

The syndrome Tuberous Sclerosis Complex (TSC) is characterized by the formation of benign tumors, referred to as hamartomas, that occur in a variety of organ types such as the brain, heart, skin, kidney, and lungs [2,13,14]. TSC mutations most frequently occur during development and displays itself in early childhood [15]. Traditionally, TSC is considered an autosomal-dominate disorder acquired through inheritance; however, approximately two thirds of TSC cases arise from sporadic germ-line mutations. Disease prevalence affects approximately 1 in 6000 live births annually and is estimated to affect roughly 1.5 million individuals worldwide [14,16,17]. TSC can arise from mutations in both the *TSC1* and *TSC2* genes; however, mutations in *TSC2* result in more severe disease related phenotypes [17–19]. Deletion, truncation, or mutation in *TSC2* can result in severe cellular outcomes [20,21]. The tremendous variability in phenotypes seen from alterations in TSC is accredited to the fact that mutations in *TSC2* span the entire length of the gene. Understanding the cellular consequences of different mutations/deletions is necessary to fully understand the overall phenotype and to identify treatment needs for individual patients.

Protein truncation test (PTT) detects changes in the DNA that can cause premature protein termination . This test has shown that truncation of the Tuberin protein is a common event in TSC patients [17]. From 18 patients analyzed by PTT six showed altered size of the Tuberin protein [22]. Single-strand conformational polymorphism (SSCP) analysis [23] has showed three TSC patients from a total of 30 presented TSC2 sequence changes predicting a truncated protein [24]. Sequencing analysis revealed a novel Tuberin truncation mutant that arises from a deletion at exon 24 in the *TSC2* gene [25]. This mutation results in a 946 amino acid protein, which is expected to retain the ability to bind Hamartin. Another Tuberin truncation protein containing only the amino acids 1-607 is still able to bind Hamartin [26]. These truncation proteins lack the C-terminal GAP domain, transcriptional activation domains, and many critical phosphorylation sites. Naturally, it can be expected that loss of these critical domains in the endogenous protein would prevent the tumour suppressor function of Tuberin and lead to uncontrolled growth resulting in tumor initiation.

Herein we analyze the effects of a clinically relevant truncation mutant of *TSC2* gene, Tuberin _[1-677],_ containing the Hamartin binding region and a portion of the CycB1 binding region. Tuberin _[1-677]_ lacks the GAP domain and many of the phosphorylation sites responsible for nutrient/growth factor regulation. Our data shows that the truncated protein has a decreased ability to retain CycB1 in the cytoplasm and thereby increases cell proliferation and mitotic onset.

## Material and Methods

### Plasmids

Full-length human pCMV-Tag2-*TSC2* mammalian expression vectors (flag-tagged) were generously supplied by J. DeClue. Human full-length CycB1-WT, -5xA, and -5xE in pCMX (GFP tagged) were generous gifts from J. Pines and have been previously described [27]. Tuberin_[1-677_] was constructed using site-directed mutagenesis by the insertion of a stop codon at the position 2050 of the *TSC2* gene inserted into the pCMV-Tag2 TSC2 vector.

### Cell Culture

HEK 293 cells (ATCC) were maintained in Dulbecco’s Modified Eagle Medium (DMEM) supplemented with 10% or 0.5 % fetal bovine serum (FBS; Thermo-Fisher) and 1% Penicillin-Streptomycin (Thermo-Fisher). Cells were incubated at 37°C in 5% CO2.

#### Antibodies

The following antibodies were utilized during Western Blotting, co-immunoprecipitation and indirect immunofluorescent analysis: Mouse α-CycB1 (monoclonal; Santa Cruz; Cat# H1809), rabbit α –Tuberin (C-4; polyclonal; Santa Cruz; Cat# 2210), mouse α-actin (monoclonal; Millipore; Cat# MAB1501), mouse α-FLAG (Sigma; Cat# F1365), goat α-mouse IgG (Sigma; Cat# A4416), goat α-rabbit IgG (Sigma; Cat#A6154), Alexa 488-conjugated α-mouse IgG (Invitrogen; Cat# 488735), Alexa 568-conjugated α-mouse IgG (Invitrogen; Cat# A-11004), rabbit α - phospho-H3 (AbCam Technologies; Cat# ab32107), rabbit α-LC3-II (Novus Biologicals; Cat# NB100-2220), rabbit α-p62/SQSTM1-(Sigma; Cat# P0067) Agarose conjugated α-FLAG primary antibody affinity gel beads (Biotool; Cat# B23101)

### Immunoprecipitation and Immunoblotting

Immunoprecipitation and immunoblotting have been previously described [9]. Briefly sub-confluent HEK 293 cells were transfected with branched or linear PEI (Polysciences, Inc.) 18 to 24 hrs after transfection the cells were lysed and incubated with CycB1 antibody overnight at 4°C and Protein G Sepharose beads (Sigma) were added to each sample for 3.5 hours at 4°C after the overnight incubation with antibody. The complexes were then washed with lysis buffer and resolved by 10% SDS-PAGE. The membrane was blotted with the indicated antisera (1:1000) followed by enhanced chemiluminescence (ECL)(Amersham). Chemiluminescence was quantified on an Alpha Innotech HD2 (Fisher) and densitometry performed using AlphaEase FC software.

### Immunofluorescent Microscopy

Cells were seeded onto glass coverslips and transfected as described above. 18 hrs following transfection, cells were fixed with 4% paraformaldehyde in phosphate-buffered saline for 2 hrs and permeabilized with 0.02% Triton X-100 for 5 min. Primary antibodies were used at 1:500 and secondary antibodies at 1:1300. Hoechst stain (Sigma) was added to the permeabilizing solution to a final concentration of 0.5μg/ml. Slides were analysed by fluorescence microscopy using Leica Microsystems or Olympus Confocal microscopy and analysed by the accompanying software.

## Results and Discussion

To analyze the effects of large deletions in Tuberin, a clinically relevant Tuberin truncated protein was created by inserting a stop codon at the position 2050 into *TSC2* gene in the pCMV-tag2 vector. The Tuberin _[1-677]_ protein contains amino acids 1-677 which comprise the Hamartin binding region and a fragment of the CycB1 binding domain (Fig 1B). To study the effects on cell proliferation the kidney cell line, HEK-293, was used given that Tuberin truncation patients present with cell overgrowth phenotypes in the kidneys [28].

Equal numbers of sub-confluent HEK-293 cells were transfected with mock, Tuberin-WT or Tuberin_[1-677]_ expression vectors. 24 hrs after transfection the number of live and dead cells were obtained via trypan blue exclusion assay and graphed as total cell numbers (Fig 2A). The proliferation assay demonstrates that despite the presence of endogenous Tuberin, Tuberin_[1-677]_ overexpression dramatically increases the number of alive cells as compared to Tuberin WT or mock control. Interestingly, protein levels of Tuberin_[1-677]_ are always significantly less than Tuberin-WT despite being driven from the same promoter (Fig 2B-C). Collectively we hypothesize that these phenotypes may occur via misregulated binding to Hamartin or CycB1, or cellular mislocalization.

**Fig 2.**
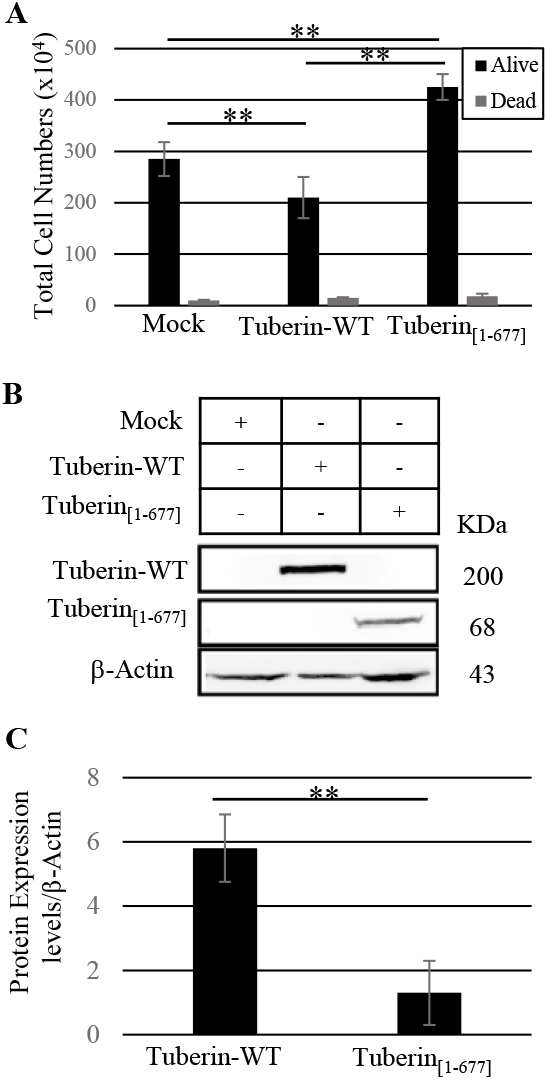
Tuberin_[1-677]_ overexpression significantly increased cell proliferation when compared to Tuberin-WT and control transfected cells. HEK-293 cells were seeded in equal numbers and transiently transfected for 24 hours with Tuberin-WT or Tuberin_[1-677]_ expression vectors. **(A)** Cell numbers counted via trypan blue exclusion assay. Total number of alive cells (black bars) and dead cells (gray bars) are depicted. **(B)** Tuberin-WT and Tuberin_[1-677]_ expression was confirmed by Western blot using Flag antibody to detect the present of the transfected proteins and B-actin was used as a loading control. **(C)** Protein expression levels were determined by densitometry and were corrected by b-actin levels. **p<0.01. Statistical significance was assessed using students’ unpaired t-test. Images and blot are representative of three experiments.

Tuberin subcellular localization is regulated, at least in part, by the Calmodulin binding-domain (CaM) which overlaps with a NLS (nuclear localization sequence) and the AKT/p90 ribosomal S6 kinase-1 and AKT phosphorylation sites located in the C-terminal of Tuberin which have been linked to nuclear localization [29,30]. To determine the subcellular localization of Tuberin_[1-677]_, the mutant was expressed in HEK-293 cells and analyzed via immunofluorescence microscopy and compared with Tuberin-WT (Fig 3; S1 Fig). Fig 3A shows confocal microscopy of both Tuberin-WT and Tuberin_[1-677]_ at 10% FBS. As expected, Tuberin-WT was found to localize primarily to the cytoplasm, however the truncated protein shows significant higher perinuclear localization when compared with the WT protein (Fig 3B). It has been demonstrated AKT phosphorylation of Tuberin at S939 increases binding to CycB1 prior to phosphorylation of CycB1 in the CRS region [9]. Reduced, but not abrogated binding between CycB1 and Tuberin occurs commitment with CycB1 phosphorylation in the CRS region and the onset of mitosis [8]. We hypothesize that the absence of these sites in the truncated protein is responsible for alterations in Tuberin localization [29].

**Fig 3.**
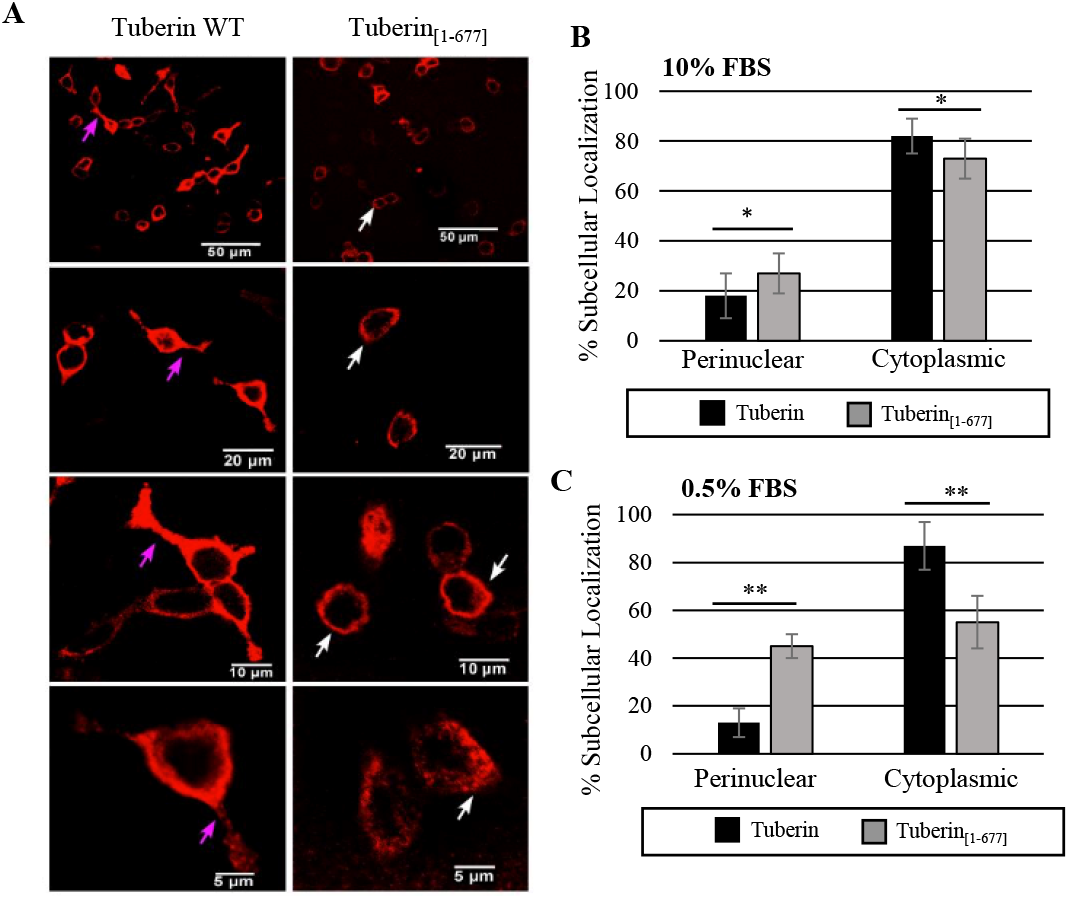
Tuberin_[1-677]_ exhibits perinuclear/nuclear localization significantly higher than Tuberin-WT. **(A)** HEK-293 cells were transiently transfected with Tuberin-WT or Tuberin_[1-677]_ expression vectors on plates containing coverslips. After 18-20 hrs the cells were subjected to either 10 % FBS (normal nutrient – panel A – S1 Fig) or 0.5% FBS (low nutrient S1 Fig) conditions for 24 hrs. Cells were collected for lysate and coverslips were subjected to immunofluorescence protocol describe in material and methods. FLAG-Texas Red (red) labelling cells expressing Tuberin-WT or Tuberin_[1-677]_. Coverslips were mounted and examined for protein localization. Pink arrows show the cytoplasmic localization of Tuberin-WT and white arrows the perinuclear/nuclear localization of Tuberin_[1-677]_. Images were done using an Olympus Confocal microscope. **(B-C)** Cell counting for Tuberin-WT and Tuberin_[1-677]_ protein localization in 10 % FSB (B) or 0.5 % FBS (C). For the full panel - S1 Fig. *p<0.05, **p<0.01. Statistical significance was assessed using a student’s unpaired t-test. Images and blot are representative of three experiments.

Reduced serum conditions (0.5% FBS) is one environmental condition prompting reduced binding between Tuberin-WT and CycB1 supporting nuclear localization and mitosis onset [9]. Hence, the cellular localization of Tuberin-WT and Tuberin_[1-677]_ were studied in serum starvation conditions (0.5% FBS) and protein localization monitored by immunofluorescence (Fig 3C; S1 Fig). Interestingly, Tuberin_[1-677]_ exhibited higher perinuclear localization, and reduced cytoplasmic localization than Tuberin-WT in both nutrient fed and starvation conditions (Fig 3B vs 3C). The increase in perinuclear accumulation during starvation for Tuberin_[1-677]_ indicates that the functional consequences for this mutation are sensitive to external stimuli.

Co-immunoprecipitation experiments were done to verify if Tuberin_[1-677]_ is able to bind CycB1 in the phosphorylated/non-phosphorylated states (Fig 4; S2 Fig). As previously demonstrated [8] and seen in (Fig 4A) Tuberin-WT affinity for CycB1 is reduced when CycB1 is modified to mimic phosphorylation on the five serine residues within the CRS (CycB1-5E), as opposed to when residues are altered to prevent phosphorylation (CycB1-5A). We have added CycB1 where S126 in the CRS region is mutated to glutamic acid and the remainder of sites are kept as a non-phosphorylatable alanine, thereby mimicking the first phosphorylation event in the CRS [27], this form is called CycB1-4A. We show that Tuberin-WT affinity for CycB1-4A is reduced as compared to CycB1-5A (Fig 4A), supporting that this first modification on CycB1 is sufficient to begin the cascade of events leading to mitotic entry. When looking at binding to the truncation mutant Tuberin_[1-677],_ which retains part of the CycB1 binding domain (Fig 1B), we note that Tuberin_[1-677]_ retains the ability to bind to CycB1 in both phosphorylated and non-phosphorylated states (Fig 4B). Previous work observed that a fragment of Tuberin (aa 610-941) was able to bind CycB1 in cells and completely *in vitro* [8]. We hypothesize based on our latest results that residues N-terminal of the CycB1 binding region in Tuberin play a role in reducing binding following phosphorylation of the CycB1-CRS region.

**Fig 4.**
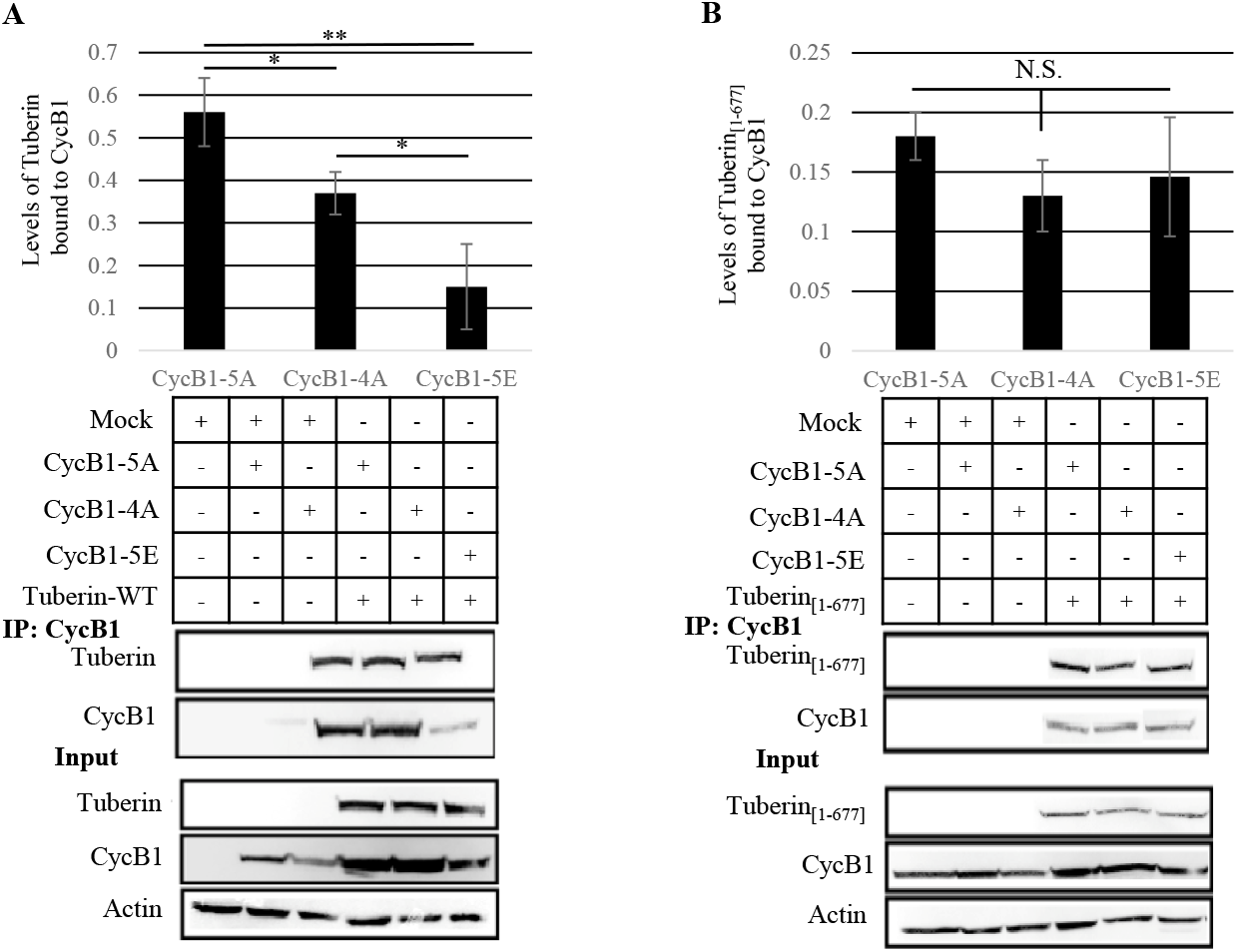
Phosphorylation state of the CycB1-CRS region alters the formation of Tuberin WT-CycB1 complex but Tuberin_[1-677]_-CycB1 lacks this level of regulation. HEK-293 cells were co-transfected using the mock (control), CycB1-5A, CycB1-4A, CycB1-5E with Tuberin-WT or Tuberin_[1-677]_ overexpression vectors. After 18 hrs the cells were collected for lysate and immunoprecipitation as described in aterial and methods using CycB1 antibody. The membrane was blotted with Flag (targeting Flag-Tuberin or Flag-Tuberin_[1-677]_), CycB1 and b-actin antibodies. **(A)** Interaction of CycB1 variants with Tuberin WT. Top panel depicts the densitometry analysis of the bands from the Western blot (bottom panel). **(B)** Interaction of the CycB1 variants with Tuberin_[1-677]._ Top panel depicts the densitometry analysis of the bands from the Western blot (bottom panel). **p<0.01, *p<0.05. (N.S.) p>0.35. Images and blots are representative of three experiments.

Tuberin delays mitotic onset by retaining CycB1 in the cytoplasm, a molecular function which is varied depending to cellular serum levels [8,9]. Immunofluorescence was performed to determine the effects of Tuberin_[1-677]_ on the subcellular localization of CycB1(Fig 5A-B; S2 Fig) and the mitotic onset (Fig 5 C-D; S3 Fig). Tuberin_[1-677]_ was found to co-localize with CycB1 as is known for Tuberin-WT (Fig 5A). CycB1 protein localization was counted as perinuclear/nuclear vs. cytoplasmic in FLAG overexpressing cells and graphed as a percentage (Fig 5B). Consistent with previous literature [8], nearly 90% of CycB1 was found to be retained in the cytoplasm when co-expressed with Tuberin-WT (Fig 5B). Interestingly, when CycB1 is overexpressed with Tuberin_[1-677]_, the subcellular localization is altered and is significantly more perinuclear/nuclear than Tuberin-WT or control cells (Fig 5B). Hence, Tuberin_[1-677]_ has lost the ability of retaining CycB1 in the cytoplasm despite still binding to it (S2 C Fig; Fig 4B), and both proteins have a perinuclear/nuclear localization. These data indicate that Tuberin_[1-677]_ lacks regulatory domains/phosphorylation sites permitting regulation by cell serum levels and retaining CycB1 in the cytoplasm, yet this truncation appears to still traffic with CycB1 to the perinuclear/nuclear space. Data is still lacking regarding the mechanistic role of this interaction at this site.

**Fig 5.**
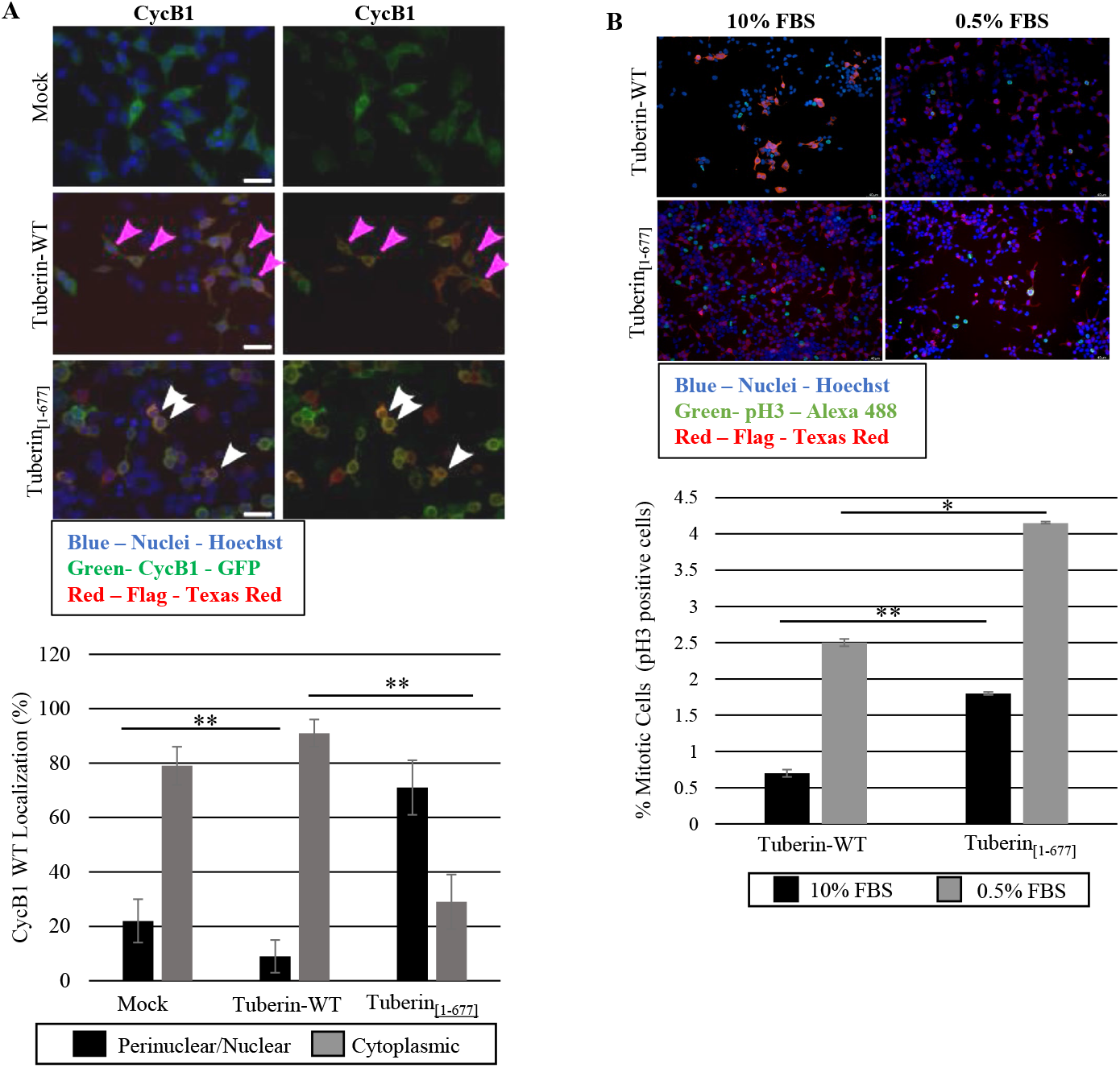
Sub-cellular localization of CycB1 and the mitotic onset are regulated differently by Tuberin-WT and Tuberin_[1-677]._ **(A)** Subcellular localization of Cyc B1 when co-transfected with Tuberin-WT or Tuberin _[1-677]._ HEK-293 cells were transiently transfected with control (mock), Tuberin-WT or Tuberin_[1-677]_ and CycB1-GFP expression vectors on plates containing coverslips. Cells were collected for lysate and coverslips were subjected to immunofluorescence protocol as described in Material and Methods. **(A – top panel)** Coverslips were mounted and examined for localization of Cyc B1-GFP (green) and Flag-Tuberin or Flag-Tuberin_[1-677]_ (red) and Hoechst (blue) was used as a nucleic marker. The figure shows the merge of the three colors. For the full panel and western blot confirming protein expression - S2 Fig. Pink arrows depict the cytoplasmic localization of CycB1 when co-transfected with Tuberin-WT and white arrows depict the perinuclear/nuclear localization of CycB1 when co-transfected with Tuberin_[1-677]_. **(A – bottom panel)** Cells were counted for their respective CycB1 locations and compared to total number of transfected cells. **(B)** pH3 was used as a marker for mitotic index in cells transfected with Tuberin or Tuberin _[1-677]_ under normal or low nutrient conditions. HEK-293 cells were transiently transfected with Tuberin-WT or Tuberin_[1-677]_ expression vectors on plates containing coverslips. **(B - top panel)** After 18-20 hrs the cells were subjected to 10% FBS (**right panel**) or 0.5% FBS (**left panel**) conditions for 24 hrs. Cells were collected for lysate and coverslips were subjected to immunofluorescence protocol as described in Material and Methods. Coverslips were mounted and examined for pH3 staining (green) and Flag-Tuberin or Flag-Tuberin_[1-677]_ (red) and Hoechst was used as a nucleic marker. The figure shows the merge of the three colors. For the full panel and western blot confirming protein expression - S3 Fig. **(B – bottom panel)** Cells were counted for pH3 staining (green) in cells expressing Tuberin or Flag-Tuberin_[1-677]_ (red) and compared to total number of transfected cells (red). Images were done using a Leica fluorescent scope. *p<0.03 **p<0.01. Statistical significance was assessed using student’s unpaired t-test. Images and blot are representative of three experiments.

The mitotic index was assessed using pH3 staining as a mitotic marker to verify if Tuberin_[1-677]_ affected levels of mitotic onset (Fig 5C-D; S3 Fig). Overexpression of Tuberin-WT indeed decreases the mitotic index when the cells are cultured in 10% FBS and no significative effect is observed at low serum condition (0.5% FBS) when comparing with the control as previously described [8,9]. Interestingly, there is a significant increase in the mitotic index when the cells are expressing Tuberin_[1-677]_ as compared to Tuberin-WT, this is seen in both 10% and 0.5% serum (Fig 5D). These results confirm the high proliferation rate obtained with cells expressing Tuberin_[1-677]_ (Fig 2A).

Tuberin mutations are a large contributing factor to misregulated cell growth pathways, supporting abnormal cell proliferation and hamartoma formation [31–35]. The results presented here indicate that clinically relevant truncation of the Tuberin_[1-677]_ protein has lost the ability to act as a tumor suppressor controlling the G2/M cell cycle transition and causes an increase in cell proliferation. Collectively, this work adds to our understanding of the function of Tuberin in regulating the sub-cellular localization of CycB1 and regulating the onset of mitosis.

These data may be relevant and important with respect to disease pathogenesis and is the first of its kind to assess a clinically relevant truncation *in vitro*. Most of the treatment available to TSC patients focus on inhibition of the mTOR protein synthesis pathways [36]. This work adds to the growing body of data supporting that it is important to understand the full scope of the biology for both WT-Tuberin regulation and that of clinical mutations and that this may result in novel treatment approaches. Additionally, this work demonstrates that imbalances in the levels of Tuberin mutants, including amplifications, can cause adverse consequences for cell biology.

## Conclusion

This is the first study to assess how this specific clinically relevant truncation of the Tuberin protein may contribute to disease manifestation. We show that elevated levels of the truncation mutant Tuberin_[1-677]_ lacking the inhibitory GAP domain retains the ability to bind and regulate the G2 cyclin CycB1. We further show that this supports increased levels of mitosis and an increase in overall cell proliferation. This work stresses that understanding the biology of different mutations/deletions in this protein is important for understanding normal and abnormal physiology.

## Acknowledgement

We thank Jiamila Maimaiti and Bob Hodge for technical assistance, Philip Habashy for his support through lab maintenance and the Imaging facility at the University of Windsor. Our work benefits from the support and input of all members of the Porter Lab, we are grateful for the diversity of contributions.

