## Supplemental Figures for "Dissecting the Roles of the Tuberin Protein in the Subcellular Localization of the G2/M Cyclin, Cyclin B1"

### Supplementary Material

**S1 Fig. Subcellular localization of Tuberin and Tuberin $_{[1-677]}$ under normal and low nutrient conditions. (A-B)** HEK-293 cells were transiently transfected with Tuberin WT or Tuberin$_{[1-677]}$ expression vectors on plates containing coverslips. After 18-20 hrs the cells were subjected to either 10 % FBS (normal nutrient – panel A) or 0.5% FBS (low nutrient – panel B) conditions for 24 hrs. Cells were collected for lysate and coverslips were subjected to immunofluorescence protocol describe in Material and Methods. FLAG-Texas Red (red) labelling cells expressing Tuberin or Tuberin$_{[1-677]}$ and Hoechst (blue) as a nuclei marker. Coverslips were mounted and examined for protein localization (A-B) and cells counting (Fig 3) for respective protein location. Pink arrows show the cytoplasmatic localization of Tuberin WT and white arrows the perinuclear/nuclear localization of Tuberin$_{[1-677]}$. Images were done using a Leica fluorescent scope. **(C)** Cell lysates were run on an SDS-PAGE gel and Western blot was performed to confirm transfection. Images and blot are representative of three experiments.

**S2 Fig. Subcellular localization of Cyc B1 when co-transfected with Tuberin or Tuberin $_{[1-677]}$. (A)** HEK-293 cells were transiently transfected with control (mock), Tuberin-WT or Tuberin$_{[1-677]}$ and CycB1-GFP expression vectors on plates containing coverslips. Cells were collected for lysate and coverslips were subjected to immunofluorescence protocol describe in Material and Methods. Coverslips were mounted and examined for localization of Cyc B1-GFP (green) and

Flag-Tuberin or Flag- Tuberin$_{[1-677]}$ (red) and Hoechst was used as a nucleic marker. Cells were counted for their respective CycB1 locations and compared to total number of transfected cells (Fig 5B). Merge 1 contains staining for Flag (red), CycB1 (green) and nuclei (blue); Merge 2 contains staining for Flag (red) and CycB1 (green) to better examine the localization of CycB1. Images were done using a Leica fluorescent scope. **(B)** Cell lysates were run on an SDS-PAGE gel and Western blot was performed to confirm transfection. Images and blot are representative of three experiments.

**S3 Fig. pH3 as a marker for mitotic index in cells transfected with Tuberin or Tuberin $_{[1-677]}$ under normal or low nutrient conditions. (A)** HEK-293 cells were transiently transfected with Tuberin-WT or Tuberin$_{[1-677]}$ expression vectors on plates containing coverslips. After 18-20 hrs the cells were subjected to 10% FBS (normal nutrients – panel A) or 0.5% FBS (low nutrients – panel B) conditions for 24 hrs. Cells were collected for lysate and coverslips were subjected to immunofluorescence protocol describe in Material and Methods.  Coverslips were mounted and examined for pH3 staining (green) and Flag-Tuberin or Flag-Tuberin$_{[1-677]}$ (red) and Hoechst was used as a nucleic marker. Cells were counted for pH3 staining (green) in cells expressing Tuberin or Flag- Tuberin$_{[1-677]}$ (red) and compared to total number of transfected cells (Fig 5D). Merge 1 contains staining for Flag (red), pH3 (green) and nuclei (blue); Merge 2 contains staining for Flag (red) and pH3 (green) to better examine pH3 staining. Images were done using a Leica fluorescent scope. **(B)** Cell lysates were run on an SDS-PAGE gel and Western blot was performed to confirm transfection. Images and blot are representative of three experiments.

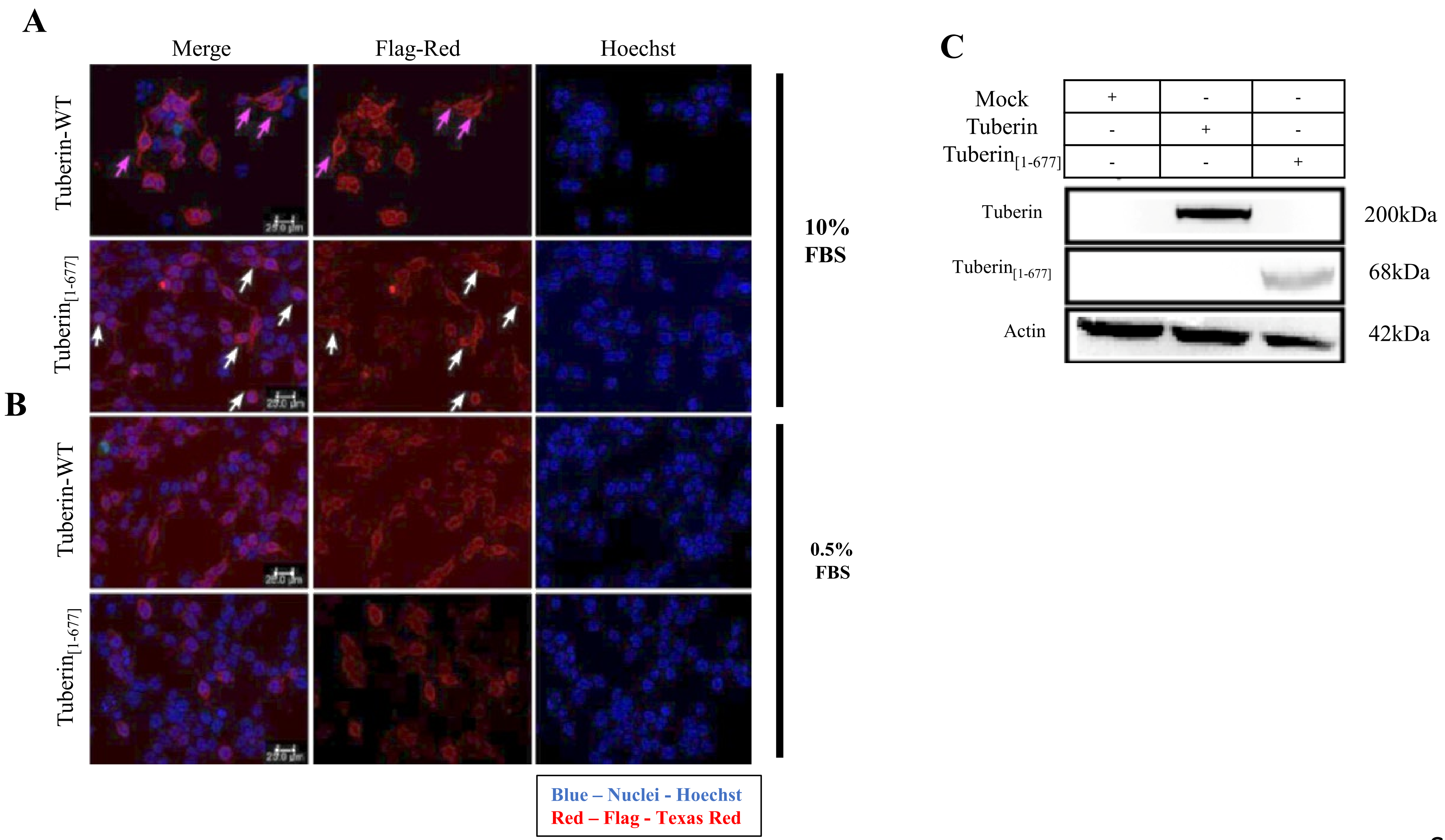
A
Merge
Flag-Red
Hoechst
Tuberin-WT
Tuberin[1-677]
10% FBS
B
Tuberin-WT
Tuberin[1-677]
0.5% FBS
Blue – Nuclei - Hoechst
Red – Flag - Texas Red
C
Mock + - -
Tuberin - + -
Tuberin[1-677] - - +
Tuberin 200kDa
Tuberin[1-677] 68kDa
Actin 42kDa

S1 Fig

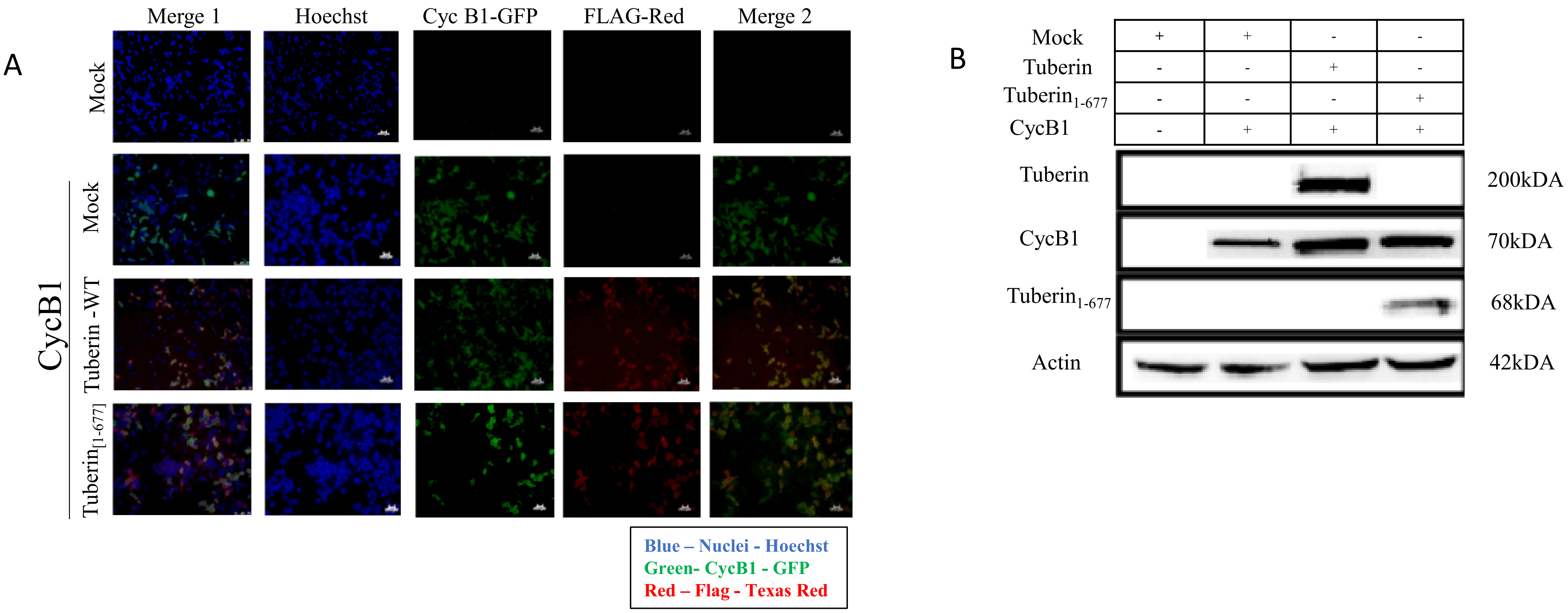
A
Merge 1
Hoechst
Cyc B1-GFP
FLAG-Red
Merge 2
Mock
CycB1
Mock
Tuberin -WT
Tuberin[1-677]
Blue – Nuclei - Hoechst
Green- CycB1 - GFP
Red – Flag - Texas Red
B
Mock + + - -
Tuberin - - + -
Tuberin1-677 - - - +
CycB1 - + + +
Tuberin
200kDA
CycB1
70kDA
Tuberin1-677
68kDA
Actin
42kDA

S2 Fig

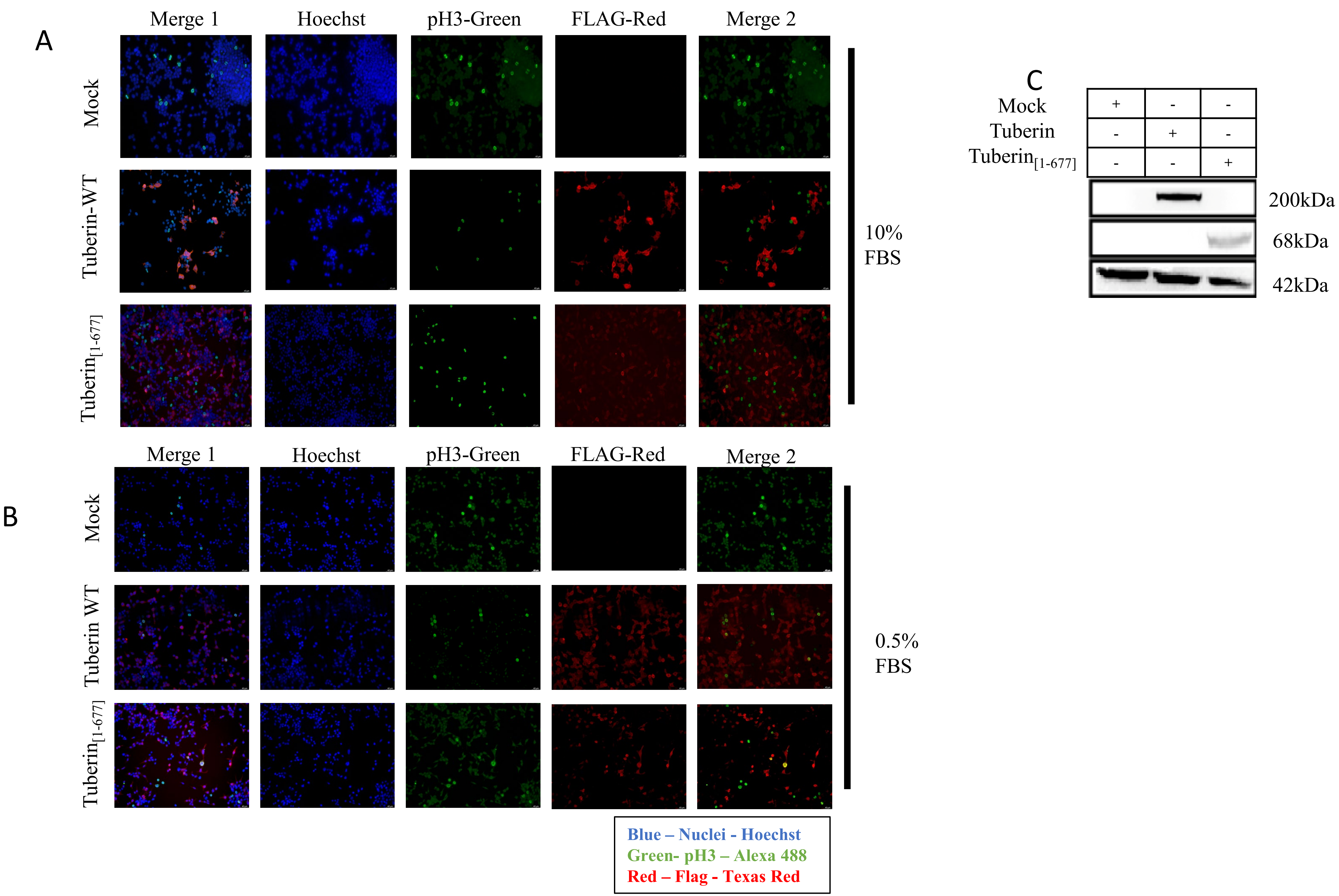
A
Merge 1
Hoechst
pH3-Green
FLAG-Red
Merge 2
Mock
Tuberin-WT
Tuberin[1-677]
10%
FBS
B
Merge 1
Hoechst
pH3-Green
FLAG-Red
Merge 2
Mock
Tuberin WT
Tuberin[1-677]
0.5%
FBS
Blue – Nuclei - Hoechst
Green- pH3 – Alexa 488
Red – Flag - Texas Red
C
Mock + - -
Tuberin - + -
Tuberin[1-677] - - +
200kDa
68kDa
42kDa

S3 Fig
